# BE3 is the major branching enzyme isoform required for amylopectin synthesis in C*hlamydomonas reinhardtii*

**DOI:** 10.1101/2023.03.08.531755

**Authors:** Adeline Courseaux, Océane George, Philippe Deschamps, Coralie Bompard, Thierry Duchêne, David Dauvillée

## Abstract

Starch-branching enzymes (BEs) are essential for starch synthesis in both plants and algae where they influence the architecture and physical properties of starch granules. Within Embryophytes, BEs are classified as type 1 and type 2 depending on their substrate preference. In this article, we report the characterization of the three BE isoforms encoded in the genome of the starch producing green algae *Chlamydomonas reinhardtii*: two type 2 BEs (BE2 and BE3) and a single type 1 BE (BE1). Using single mutant strains, we analyzed the consequences of the lack of each isoform on both transitory and storage starches. The transferred glucan substrate and the chain length specificities of each isoform were also determined. We show that only BE2 and BE3 isoforms are involved in starch synthesis and that, although both isoforms possess similar enzymatic properties, BE3 is critical for both transitory and storage starch metabolism. Finally, we propose putative explanations for the strong phenotype differences evidenced between the *C. reinhardtii be2* and *be3* mutants, including functional redundancy, enzymatic regulation or alterations in the composition of multimeric enzyme complexes.

## Introduction

Viridiplantae, as well as various algae, store carbon into an insoluble and semi-crystalline homopolymer of glucose called starch (Pérez and Bertoft, 2010). This polysaccharide is composed of linear α-1,4 linked glucans ramified through α-1,6 bonds, also called branching points. Starch granules usually contain two fractions corresponding to polysaccharides of distinct architectures. The minor fraction, called amylose, accounts for 10 to 25% of wild-type starches and is mostly composed of long linear glucans. Alternatively, the majority of the wild-type starch granule dry weight (75-90%) is made of a moderately branched polysaccharide, called amylopectin, containing around 5% of α-1,6 linkages with a degree of polymerization (DP) of approximately 10^5^ (Manners, 1989). Amylopectin synthesis in Plantae is a complex mechanism that requires the concerted action of many enzymes, including several isoforms of starch synthases (SS; E.C. 2.4.1.21), branching enzymes (BE; E.C. 2.4.1.18) and isoamylase-type debranching enzymes (DBE; EC. 3.2.1.68). That biosynthetic pathway leads to the formation of highly ordered amylopectin molecules which are responsible for the physico-chemical properties of the polysaccharide such as its insolubility and its semi-crystalline nature. Starch synthases are in charge of the building of the linear glucose chains, branching enzymes catalyze the hydrolytic cleavage and transfer of an α-1,4 bond within the glucan chain and give rise to α-1,6-branch linkages. The branched polysaccharide obtained requires a trimming process catalyzed by isoamylases to achieve the fine structure of amylopectin (Ball et al., 1996). Branching enzymes belong to the α-amylase superfamily (GH13) in the CAZy database (Lombard et al., 2014). They share a common three-domains structure composed of a N-terminal domain containing a carbohydrate binding module (CBM48), a central catalytic domain found in the GH13 family members and a carboxy-terminal β-domain found in many α-amylases. Several BE isoforms are found in plants and algae and classified as class 1 or class 2 enzymes based on sequence similarity (Boyer and Preiss, 1978). A single isoform of class 1 is usually detected in plants, but may be missing (*e.g.* in Brassicaceae). A single isoform of type 2 BE also generally occurs, like in potato and pea. Conversely, two type 2 BE isoforms are found in cereals, like in maize, rice or barley where BEIIa and BEIIb display distinct expression patterns, the latter being exclusively found in the grain (Gao et al., 1997; Sun et al., 1998; Yamanouchi and Nakamura, 1992) while BEIIa is expressed in all tissues.

Many studies performed on plant BEs revealed the absolute requirement of a branching activity for amylopectin synthesis. In general, specific defects for class 1 BE cause mild or no phenotype on starch structure or accumulation in maize, rice or wheat endosperms questioning the role of this BE family in starch metabolism (Regina et al., 2004; Satoh et al., 2003; Xia et al., 2011). On the other hand, the complete loss of starch branching activity in Arabidopsis mutant plants that are disrupted for both type 2 BEs leads to the disappearance of starch (Dumez et al., 2006). Additionally, partial defects in class 2 BEs (BEIIa or BEIIb) lead to the strong *amylose extender* (*ae*) phenotype and has been described in many plant mutants including the *beII* mutants of pea, the *beIIb* mutants of rice and maize, and in lines silenced for BEIIa of durum wheat (Bhattacharyya et al., 1990; Hedman and Boyer, 1982; Mizuno et al., 1993; Sestili et al., 2010). The *ae* phenotype consists of an increase of the amylose content as well as a modified amylopectin structure containing longer internal and external chains. However, in maize *beIIa* mutants, the endosperm starch displays a wild-type phenotype but leaf starch is substituted by a low molecular weight poorly branched polysaccharide. In this specific case, the BE1 isoform may be involved in the synthesis of this residual polysaccharide (Yandeau-Nelson et al., 2011). Altogether, these studies revealed the prime importance of branching enzymes in determining the correct amylopectin structure and the physico-chemical properties of the produced starch.

The unicellular green alga *Chlamydomonas reinhardtii* is a long-time established model for forward genetics. This photosynthetic microbial organism offers several advantages including: a mainly haplontic lifestyle, a short generation time compared to land plants, several whole genome sequences assembly and annotation, efficient molecular tools and resources, and a controlled sexual cycle allowing co-segregation and complementation analyses. Chlamydomonas has been extensively used as a model organism for studying chloroplast biology, photosynthesis, or flagellar assembly (Harris, 2001). Moreover, Chlamydomonas produces starch granules similar to those found in land plants both morphologically and in composition and structure (Buléon et al., 1997), and starch metabolism can be tuned depending on growth conditions (Libessart et al., 1995). In photosynthetically active cells, photosynthetic (or transitory) starch is produced as a sheath around the pyrenoid, and presents similar properties to the one accumulated in land plants source organs (like leafs). After a few days under restricted growth conditions, transitory starch is replaced by a storage-like polysaccharide, accumulated in large quantities in the chloroplast stroma, and harboring the characteristics of land plants endosperm storage starch (Delrue et al., 1992; Fontaine et al., 1993; Libessart et al., 1995; Maddelein et al., 1994). This behavior emulates the tissue-specific starch metabolism of plants, and switching between growth conditions enables the deciphering of the modulations, from the same set of enzymatic functions, that allows either the synthesis of transitory starch or the use of starch for long term carbon storage. The detailed cellular mechanisms underlying the switch between transitory to storage starch accumulation in Chlamydomonas is still not known, although one related gene function was recently identified (Findinier et al., 2019). Phenotype screening of mutants with potential alteration in starch metabolism have been historically conducted by exploiting the specific interaction of molecular iodine with α-glucans. Chlamydomonas colonies stained with iodine display modification from the wild-type dark blue into a range of colors evidencing alterations in starch structure or content. These screens have led to the identification of many gene encoding functions involved in starch metabolism (Hicks et al., 2001; Tunçay et al., 2013). The early availability of the annotated Chlamydomonas nuclear genome sequence has been decisive for gene identification in “forward genetics” strategies. (Merchant et al., 2007). However, mutants for several genes that are strongly suspected to be involved in starch metabolism were never isolated through this approach, probably because their phenotypes are masked by redundancy.

In this work, we studied several insertion mutant lines defective for each of the three starch branching enzyme isoforms encoded in the genome of *Chlamydomonas reinhardtii*. We characterized the structural and metabolic consequences of these alterations for both transitory and storage starch. The loss of the class 1 branching enzyme isoform (BE1) does not affect starch anabolism at all. Disruption of the genes encoding one of the two type 2 BEs (BE2 or BE3) leads to a modified starch structure. Our data suggest a preponderant role for BE3 in starch metabolism that can’t be compensated by any of the two other isoforms. We also purified recombinant proteins of the three BE isoforms and determined the substrate preferences and the lengths of the α-glucan chains transferred by each enzyme. Possible explanations for the phenotype differences that are observed between the *be2* and *be3* mutants are also discussed.

## Results

### Survey and structure analysis of Chlamydomonas branching enzymes

The nuclear genome of *Chlamydomonas reinhardtii* contains three genes encoding starch branching enzymes: *Cre06.g289850*, *Cre06.g270100* and *Cre10.g444700;* annotated BE1, BE2, and BE3 respectively. The corresponding proteins all harbor the structural characteristics of starch branching enzymes including: (1) a N-terminal CBM48 carbohydrate-binding domain; (2) a central (β/α)_8_-barrel catalytic domain containing four highly conserved regions, which is a characteristic feature of the GH13 family (Stam et al., 2006); (3) a C-terminal domain involved in controlling enzyme catalysis as well as in determining glucan substrate preference (Kuriki et al., 1997; Hong and Preiss, 2000). They also contain the aspartic-acid/glutamic-acid/aspartic-acid triad, which is essential for catalytic activity (Fig.1). Additionally, chloroP detects a N-terminal chloroplast targeting transit peptide in the protein sequence of all three isoforms (Emanuelsson et al., 1999). BE2 and BE3 share 80 % sequence similarity and 71 % identity whereas these values fall to 67 % similarity and respectively 54 % and 53 % identity when compared to BE1. A gene phylogeny of the three Chlamydomonas BE proteins was inferred to determine how they relate to their homologs in algae and plants (Fig. 2 and Fig. S2). All BE in Viridiplantae share a common ancestor gene with other Archaeplastida (Deschamps et al., 2008), allowing us to use Rhodophyceae as an outgroup. Our gene tree indicates that type 1 and 2 BE are paralogs derived from a duplication in the ancestor of all Viridiplantae. The high level of conservation of these two paralogs across the phylum is evidence of their metabolic importance. Chlamydomonas BE1 is related to all type 1 BE which are described to primarily transfer longer chains. No additional duplications were detected within the BE1 clade. Alternatively, several independent events of duplication can be observed within type 2 BE: in Chlamydomonadales, in Poaceae and in Brassicaceae. Type 2 BE of land plants are responsible for transferring shorter chains and we may infer that this would also be the case for Chlamydomonas type 2 BE. However, Chlamydomonas BE2 and BE3 don’t share a specific duplication ancestry with BEIIa and BEIIb of Oryza and neither with BE2.1 and BE2.2 of Arabidopsis. Thus, if all those paralogs should have similar specific sub-functional traits, it would be the result of convergent evolution. In order to have a closer look at type 1 and 2 shared structural properties, BEs protein sequences of *Chlamydomonas reinhardtii*, *Oryza sativa*, *Solanum tuberosum* and *Zea Mays* were compared in a multiple alignment and by inferring their secondary structure (Fig. 2A-C). A recent study proposed that the specificity of plant BE enzymes regarding the length of the glucan they transfer is conditioned by the composition of two specific loops in their sequence (Gavgani et al., 2022). We used Alphafold to predict the structure of the three Chlamydomonas isoforms (Fig. S1). Interestingly, the two aforementioned loops are found in the Chlamydomonas BEs protein sequences. In particular, a specific insertion found in rice BEIIb-type isoform is also observed in the second loop (loop 541, Fig. 1C) of both Chlamydomonas BE2 and BE3. Protein structure predictions using AlphaFold confirmed these similarities between the loops of the rice and the Chlamydomonas enzymes (Fig. 2) suggesting once again that BE1 resembles a type 1 BE while both BE2 and BE3 are closely related to type 2 BEs. In land plants, starch branching enzyme isoforms are regulated by protein phosphorylation which regulates the association of BEs in multienzyme protein complexes (Tetlow et al., 2008; Hennen-Bierwagen et al., 2008, 2009). Three serine residues are phosphorylated in the plant class II BEs, Ser^286^ and Ser^297^ and Ser^649^. Sequence alignment revealed that the three putative phosphorylation sites are found in Chlamydomonas BE2 while the BE3 isoform lacks the first serine (Fig. 2C).

**Figure 1:**
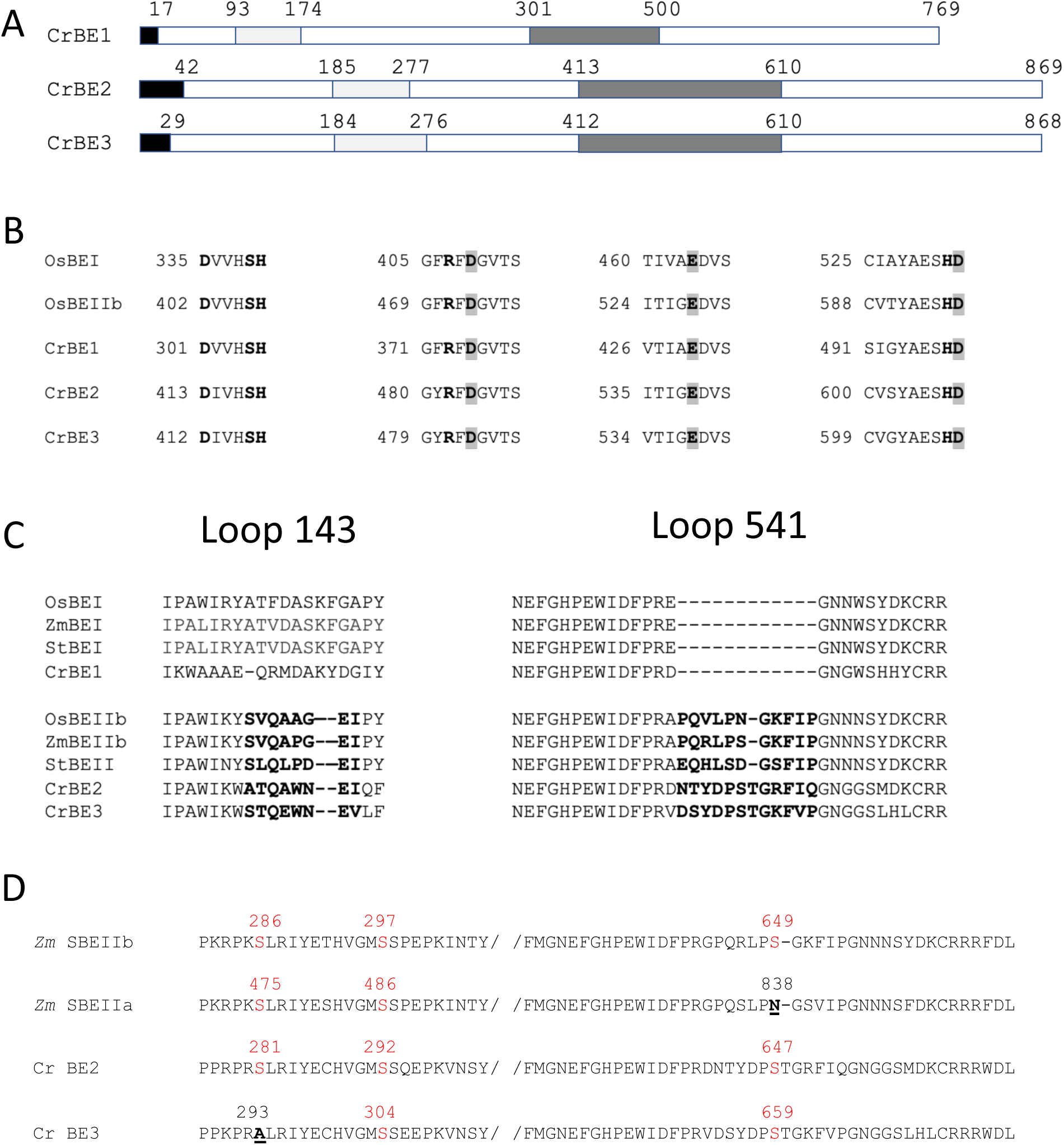
Chlamydomonas starch branching enzymes organization. (A) Functional domain organization of Chlamydomonas starch branching enzyme isoforms showing the putative transit peptide in black, the carbohydrate binding module CBM 48 in the N-terminal part (in light grey), and the central catalytic core containing four highly conserved regions (in dark grey). (B) Sequence alignment of the four conserved regions contained in the catalytic core found in the Chlamydomonas BE isoforms and the rice BEI and BEIIb. The catalytic triad “DED” is enlightened in grey and the invariant amino acids residues are in bold. (C) Sequence alignment between the two loops (143 and 541) which were shown to distinguish land plant BEI and BEIIb isoforms and the corresponding region in Chlamydomonas BE isoforms (Gavgani et al., 2021). (D) Sequence alignment displaying the phosphorylation sites found in maize SBEIIb and SBEIIa and the Chlamydomonas BE2 and BE3 isoforms. The serine residues are displayed in red, when missing the amino acid residue replacing the serine residue is underlined and displayed in bold. Numbering corresponds to the complete peptide sequence of the indicated protein on each figure panel. Os BEI: B7EAH2, Os BEIIb: B3VDJ4, Zm BEI: Q7DNA5, St BEI: P30924, St BEII: Q9XGA6, Zm SBEIIa: O24421, Zm SBEIIb: Q08047.

**Figure 2:**
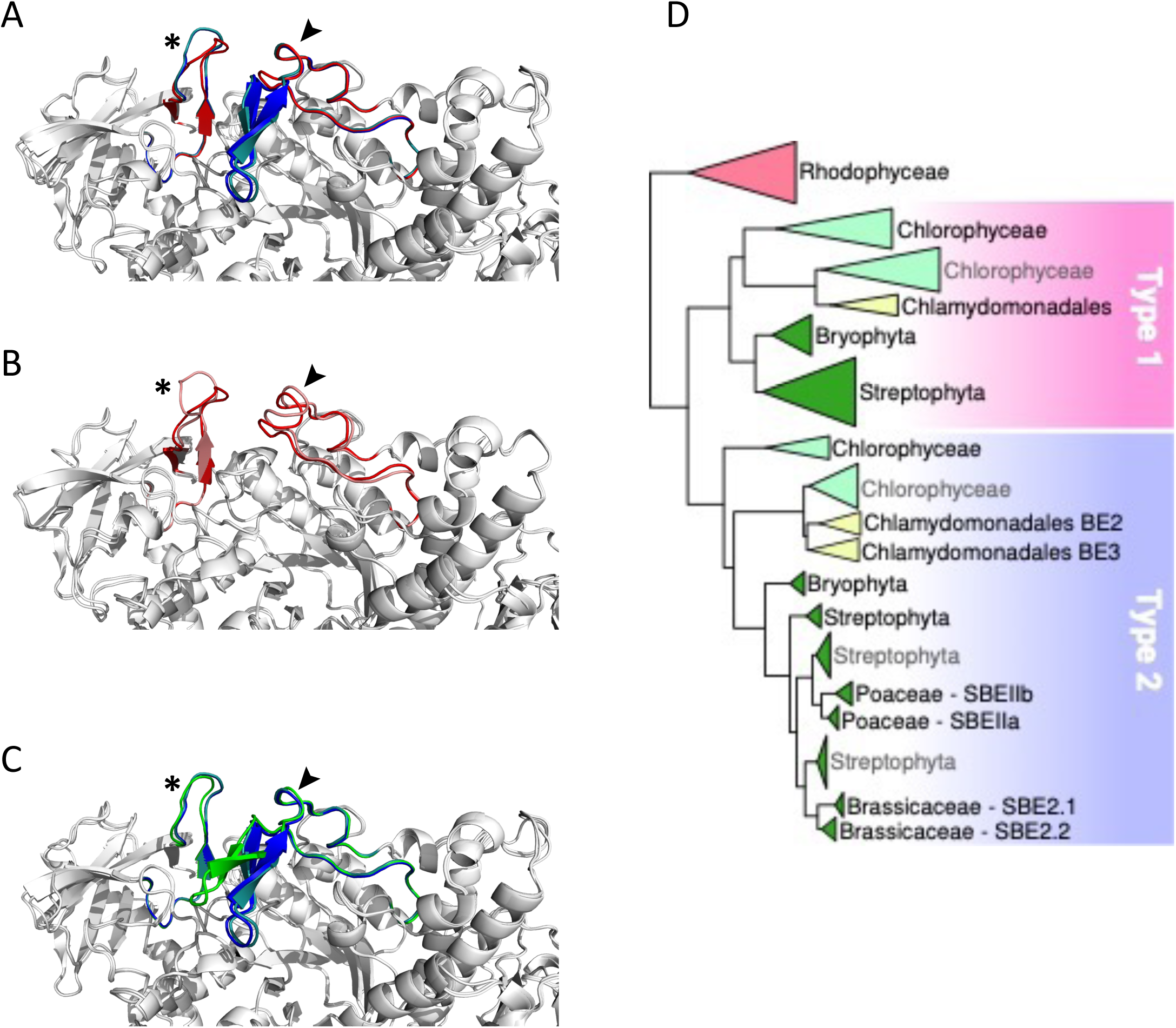
Comparison of functionally important loops in Branching Enzymes from *Chlamydomonas reinhardtii* (Cr) and *Oryza sativ*a (Os) and schematic phylogenetic tree of starch branching enzymes. Panels A to C display close up on the conserved specific loops as described in Gavgani et al. (2022) on overlays of the different enzymes represented as white cartoon. Loops 143 and 541 are identified by a star and an arrow respectively. (A) overlay of Alphafold structures of the three CrBEs, CrBE1 (red), CrBE2 (navy blue) and CrBE3 (light blue). (B) overlay of the Alphafold structure of CrBE1 in red and the crystal structure of OsBE1 in salmon (pdb code 7ML5, Galvgani et al., 2022) (C) overlay of Alphafold structures of CrBE2 (navy blue), CrBE3 (light blue) and OsBE2 (green). (D) Schematic phylogenetic tree of starch branching enzymes in algae and plants resuming the detailed tree presented in figure S2. Please refer to M&M for details on tree reconstruction methodology. Branching enzymes of Rhodophyceae were used to root the tree. Type 1 and type 2 BE are highlighted by a pink and a blue box respectively.

### Selection and characterization of branching enzymes disrupted mutants of Chlamydomonas

For each of the three branching enzyme genes described above, two independent insertion lines were selected from the Chlamydomonas CLiP mutant library (Li et al., 2019) (see M&M for accession numbers). Insertion tags in exons were favored to optimize chances of obtaining strains deficient for each gene’s corresponding activity. PCR amplifications of genomic DNA purified from each strain were performed using primers specific to flanking regions of the exogenous DNA insertion site (Table S1). All mutant strains proved to have a specific interruption in the expected exon (Fig. S3), leading to the loss of expression of the corresponding genes (Fig. 3A).

**Figure 3:**
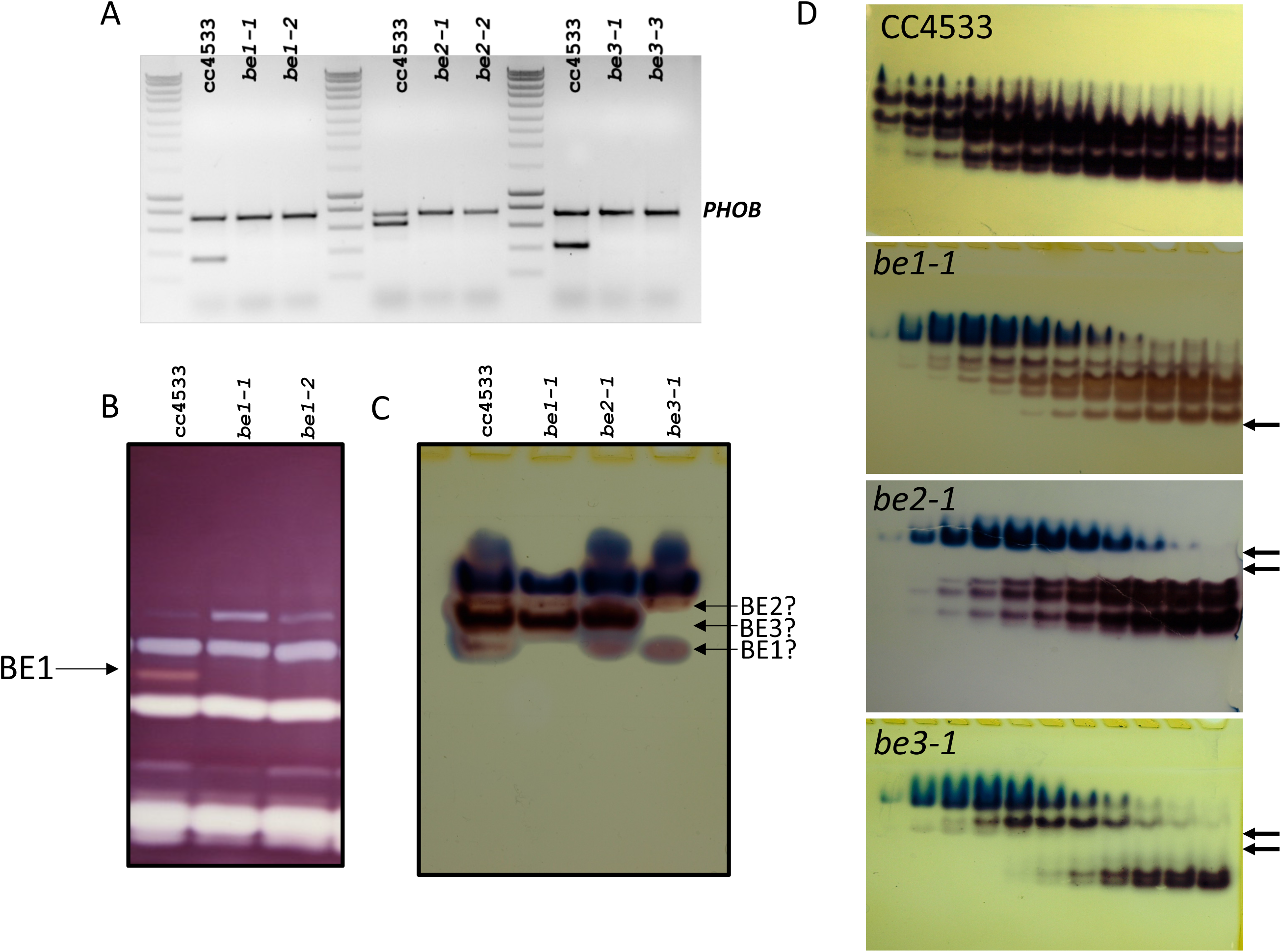
Characterization of the missing activities in the Chlamydomonas mutants defective for BE isoforms. (A) Expression of the *BE* mRNA in the wild-type (CC5325) and the mutant alleles. Total RNA (500 ng) extracted from the WT and the mutant strains were used to amplify specific fragments of the BE genes by RT-PCR. A 736 bp fragment of the starch phosphorylase gene (*PHOB*) was amplified as a control. Molecular weight marker: Smartladder (Eurogentec). The primers sequence and the expected size of the amplicons can be found in supplemental table 1. (B) Denatured crude extracts (25 µg of protein) from the wild-type and *be* mutant strains were loaded on denaturing polyacrylamide gels containing 0.3 % soluble potato starch (Mouille et al., 1996). The proteins were renatured after electrophoresis and incubated overnight. Starch hydrolases activities were revealed by staining the gel with Lugol. The 84-kD pink-staining band was previously proven to correspond to BE1 (Tunçay et al., 2013). (C) Crude extracts from the wild type and *be* mutant strains were analyzed on native PAGE (8 % acrylamide) and the gels were incubated overnight in the presence of glucose-1-phosphate and rabbit muscle phosphorylase a. Starch branching enzyme activities appear as reddish bands while blueish bands correspond to the endogenous Chlamydomonas starch phosphorylase activities. A possible assignment of each Chlamydomonas isoform is proposed on the right of the figure. (D) Anion exchange chromatography (Hi-trap Q) performed on 100 mg of proteins from the wild-type and the mutant strains. Proteins were eluted by a linear gradient of NaCl (0 to 300 mM in 90 min, 1.5 ml fractions) and aliquot (30 µl) of each even fraction from fraction 10 to 32 were used to visualize BE activities on native zymograms as in (C). Arrows indicate the missing activities in each mutant strain.

Biochemical analyses were then conducted to determine if those strains are indeed devoid of the relevant enzymatic activities. On zymograms revealing alpha-glucan hydrolytic activities and performed under denaturing conditions, only the BE1 isoform could be detected. This activity modifies the starch matrix contained in the gel and is revealed by the appearance of a pink band after iodine staining (Fig. 3B). The specific lack of this BE isoform was confirmed for both *be1-1* and *be1-2* mutant alleles (Fig. 3B). Unfortunately, under these experimental conditions, the other putative branching activities cannot be observed. This may be due to the denaturation/renaturation procedure, or by confounding enzymatic activities acting on the same region of the zymogram. Indeed, Chlamydomonas BE2 and BE3 mature proteins have an estimated molecular weight of about 90 kDa, which is similar to Chlamydomonas isoamylases, possibly resulting in the masking of the branching activities by the debranching enzymes. In order to visualize BE2 and BE3, specific zymograms were performed under native conditions. Several activities were detected in the wild type strain, suggesting that the branching enzyme isoforms may be organized into protein complexes (Fig. 3C). However, it is worth noting that different enzymatic extracts from the same strain displayed activity profiles which varied greatly in terms of band intensities, excluding the use of this technique to highlight the specific lack of one isoform in the mutants. Furthermore, interpretation of these gels was complicated by the presence of endogenous starch phosphorylase activities forming blue bands at the top of the gels (Fig. 3C). To overcome this difficulty, the BE containing protein complexes were separated using anion-exchange chromatography before performing zymograms in native conditions (Fig. 3D). With this procedure, clear differences between wild-type and mutant strains were evidenced. For each genotype, we were able to attribute specific activity bands and confirm the specific loss of some of them in our mutants (Fig. 3D). In the *be1* mutant background, only the low intensity fast migrating band specifically disappeared. Four additional bands corresponding to BE activity were detected and attributed to either BE2 or BE3. For each mutant, two of them specifically disappeared (Fig. 3D). The *be2* mutation led to the disappearance of the two slowest migrating activity bands, whereas protein extracts from *be3* deficient strains no longer contained the two activity bands detected in the middle of the gel (Fig. 3D). Finally, one BE activity band migrating just above BE1 was detected in all mutant extracts. This band is either revealing an additional BE isoform that was not identified yet in the genome, or may constitute an alternative multimeric complex, possibly composed of BE2 and BE3 units.

### Phenotype, starch content and structure analysis of branching enzyme’s single mutants

We studied the consequences of the loss of each BE isoform on starch synthesis in Chlamydomonas. The loss of one starch branching enzyme isoform did not affect the growth of mutant strains (Fig. S4). We analyzed starch content and structure produced by cells collected, either in exponential growth conditions, or after 5 days under nitrogen starvation. For the sake of clarity, we will thereafter refer to starch collected in each condition as transitory starch and storage starch respectively.

The amount of transitory and storage starch accumulated by the wild-type and in mutant strains were calculated from 5 independent experiments for each strain and displayed as the means ± SD in Table 1. Under both growth conditions, *be1-1* and *be1-2* mutants accumulated similar amounts compared to the wild-type reference strain. The *be3* mutants accumulated 20 % less transitory starch compared to the wild-type, whereas strains disrupted at the *BE2* locus surprisingly accumulated slightly more (120 %) (Table 1). Alternatively, both *be2* and *be3* mutants showed a slight decrease in storage starch accumulation compared to the reference strain. (Table 1).

**Table 1:** Phenotype of wild-type and mutant strains during transitory (+N) or storage (- N) starch synthesis. The results are the average of at least 5 independent experiments performed on independent samples and are displayed as means ± SD. The amounts of starch expressed in µg per 10^6^ cells were obtained after purification of the insoluble polysaccharide and were measured by the standard amyloglucosidase assay. The amylose ratio and the λmax of amylopectin were obtained after purification through size exclusion chromatography on sepharose CL-2B columns. Statically significant differences with the wild-type values are indicated by a star (p<0,05).

| STRAINS | +N |  |  | -N |  |  |
| --- | --- | --- | --- | --- | --- | --- |
| | Starch | $\lambda_{\max}$ | Am% | Starch | $\lambda_{\max}$ | Am% |
| CC4533 | 6,2 ± 1,3 | 579 ± 1 | 7,5 ± 1,9 | 63 ± 4 | 558 ± 5 | 35,3 ± 6,1 |
| *be1-1* | 5,9 ± 2,1 | 581 ± 3 | 7,3 ± 0,5 | 68 ± 7 | 556 ± 6 | 35,2 ± 5,2 |
| *be1-2* | 5,7 ± 1,2 | 583 ± 1 | 7,8 ± 2,2 | 64 ± 5 | 557 ± 4 | 38,9 ± 6,2 |
| *be2-1* | 7,3 ± 0,9* | 553 ± 3* | 9,5 ± 2,7* | 49 ± 8* | 549 ± 4* | 26,5 ± 2,5* |
| *be2-2* | 7,5 ± 1,3* | 555 ± 4* | 11,6 ± 1,4* | 51 ± 4* | 548 ± 5* | 28,8 ± 2,1* |
| *be3-1* | 5,5 ± 1,3* | 580 ± 4 | 16,1 ± 2,3* | 46 ± 11* | 569 ± 3* | 46,2 ± 4,5* |
| *be3-3* | 4,9 ± 1,6* | 578 ± 5 | 15,8 ± 1,2* | 53 ± 6* | 571 ± 6* | 48,1 ± 5,8* |

Amylose and amylopectin were fractionated using CL2B gel permeation chromatography performed on the insoluble polysaccharide produced by the wild-type strain and the *be1*, *be2*, and *be3* mutants. A representative profile obtained for the polysaccharides produced by each mutant is displayed in figure 4 for both transitory (Fig. 4A to 4D) and storage starch (Fig. 4E to 4H). These analyses allowed us to identify modifications in the amylopectin/amylose ratio as well as in the structure of amylopectin (evidenced by a shift in the λmax value of the amylopectin/iodine complex) (Table 1).

**Figure 4:**
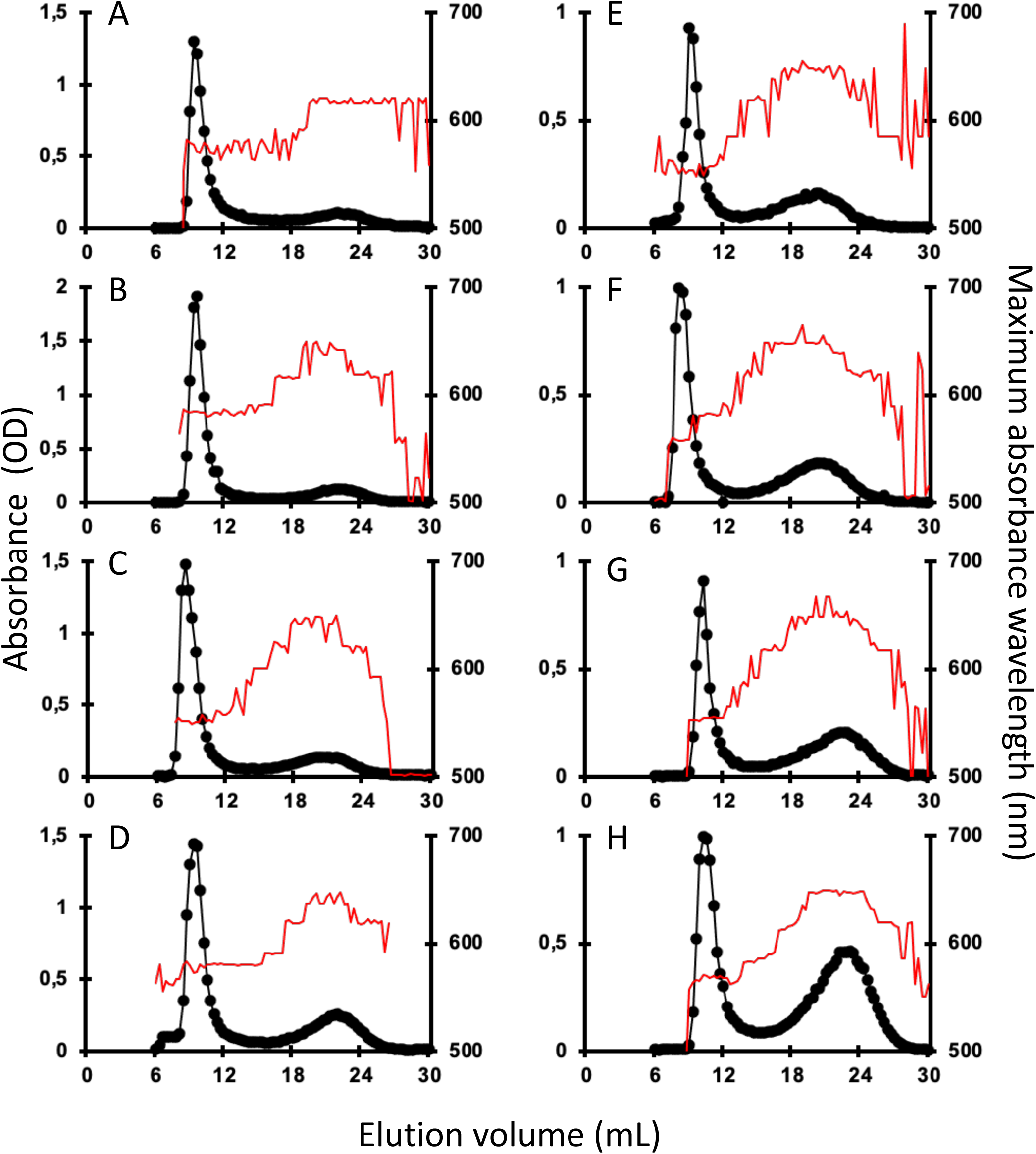
Separation of amylopectin and amylose by CL2B-gel permeation chromatography. The optical density (•) was determined for each 300 µl fraction at the maximum wavelength (indicated by the red thin line). All samples were run on the same column setup described by Delrue et al. (1992). Starches were extracted from exponentially growing cells (A to D) or nitrogen-starved cultures (E to H) from the wild-type reference strain CC5325 (A and E), the *be1-1* (B and F), the *be2-1* (C and G), and *be3-1* (D and H) were analyzed. The chromatograms shown are from one representative experiment chosen from at least five independent experiments; the same characteristics were observed in each of the second mutant alleles (*be1-2, be2-2*, and *be3-3*). Quantification of amylose and amylopectin ratios was obtained by pooling amylopectin and amylose fractions and measuring the amount of glucose through the standard amyloglucosidase assay (Table 1). The amylopectin λmax values displayed in table 1 were measured after mixing an aliquot (80 µl) of the amylopectin pool with iodine.

Transitory starch produced by both the wild-type and the *be1* mutant strains contain comparably low relative amounts of amylose. Conversely, the amylose content is increased 1.3- and 1.5-fold in *be2* mutants and 2.4- and 2.1-fold in *be3-1* and *be3-3* strains respectively (Table 1). Moreover, storage starch in *be3-1* and *be3-3* mutants is also enriched in amylose compared to the reference strain (46 and 48% of amylose respectively compared to 35 % in the wild-type). Surprisingly, both *be2* mutants display an unexpected phenotype with a decrease in amylose relative content (26 and 29 % for *be2-1* and *be2-2* respectively). The structure of amylopectin in transitory starch was globally unchanged for all mutants compared to the reference strain (λmax≃580 nm), with the noticeable exception of the *be2* mutant where the λmax of amylopectin dropped to 555 nm (Table 1). This lower value is comparable to what is usually observed for wild-type amylopectin in storage starch (λmax=560 nm). The amylopectin molecules from storage starches produced by both the wild-type and the *be1* mutants display similar λmax values. A moderate decrease of amylopectin λmax was systematically recorded in storage starch of the *be2* mutants, though these differences were not statistically significant (Table 1). Finally, the two *be3* mutants produced an amylopectin with an increased λmax value (570 nm) which is usually described in land pants BE mutants.

In order to identify the reasons for these changes, we studied the chain composition of each polysaccharide in more detail. Five milligrams of starch purified from the wild-type and the mutant strains were debranched using a cocktail of isoamylase and pullulanase and the resulting linear chains were separated on a TSK-HW50 gel permeation column. An aliquot of each fraction was then mixed with iodine to determine the maximum absorbance and the corresponding λmax value allowing to determine the average length of the linear chains composing each fraction (Fig. 5). The wild-type starch profile (Fig. 5) was composed of two peaks, the first corresponding to the amylose subfraction and to the long chains of amylopectin (degree of polymerization (DP) greater than 80 glucose residues) while the second peak contained chain sizes of smaller DP. The profiles obtained for the transitory (Fig. 5A) and storage starches (Fig. 5B) were similar except for the intensity of the first peak which corresponds mainly to the amylose chains. Again, the profiles obtained for the *be1* mutants were almost identical to those obtained for the wild-type reference. Transitory starch (Fig. 5A) produced by the *be2* mutants contains fewer long chains in amylopectin (DP > 80). The opposite observation was made for starch produced by *be3* mutants, which displayed an enrichment for this population of long chains compared to the wild-type reference (Fig. 5A). The same trends were observed for storage starches of *be3* mutant strains (Fig. 5B) while an enrichment in chains with DP composed of less than 70 glucose residues was assessed in the *be2* mutants.

**Figure 5:**
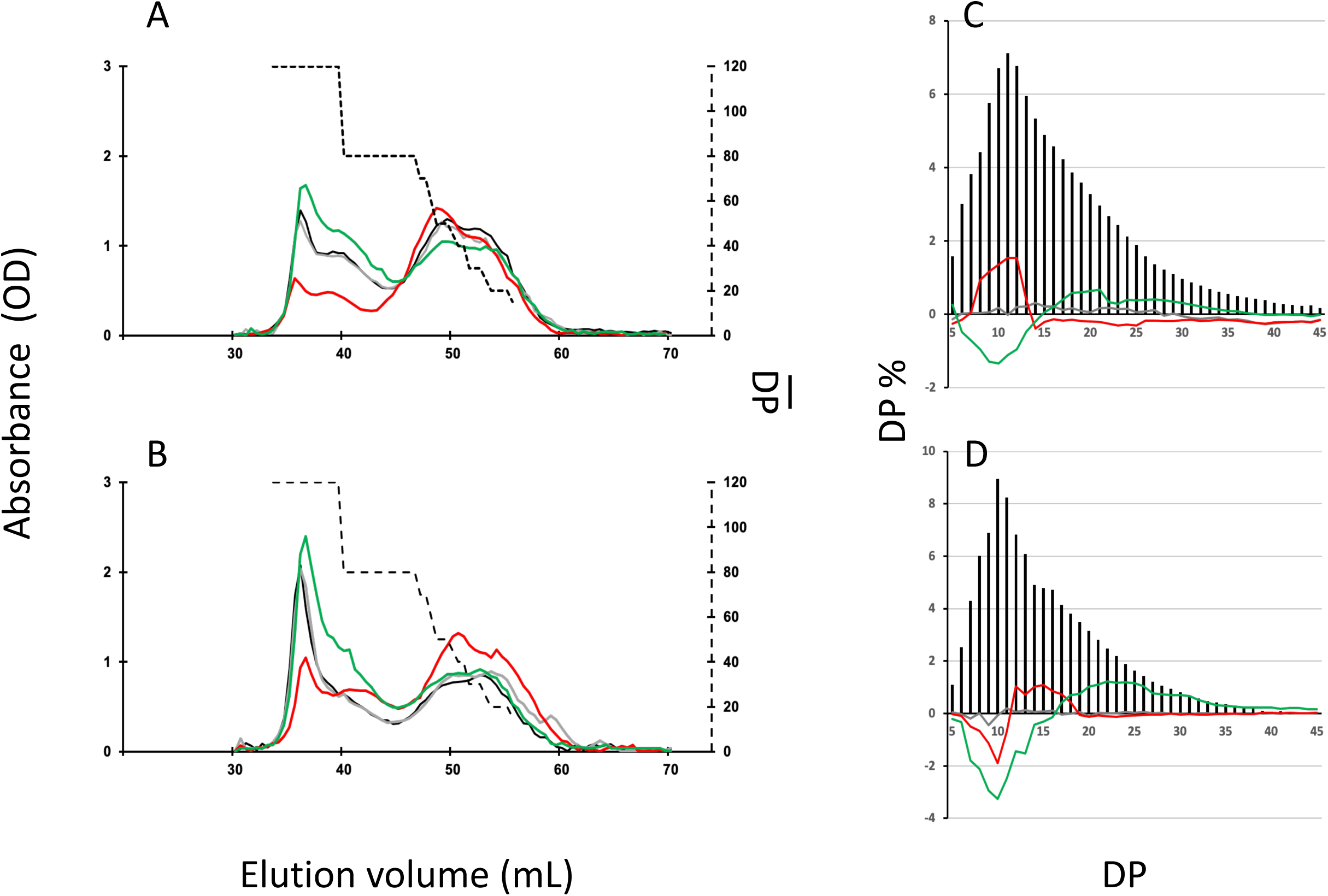
Separation of starch and amylopectin constitutive chains after enzymatic debranching. (A) and (B), Starch (10 mg) extracted from the wild-type and the mutant strains were debranched by a cocktail of isoamylase and pullulanase (see methods) and the debranched chains were separated onto a TSK-HW50 gel permeation chromatography equilibrated in 10 % DMSO at a flow rate of 12 ml/h. Absorbance at the maximum wavelength of the iodine/polysaccharide was determined for each fraction. Chromatograms from one representative experiment performed on the wild-type (black), the *be1-1* (grey), the *be2-1* (red), and the *be3-1* (green) mutants are displayed. These analyses were performed for both transitory starch (exponentially growing cells, A) and storage starch (nitrogen starvation, B). The average DP (broken line, right y-axis) was generated by using the λmax values of the debranched glucans as internal standards according to Banks et al. (1971). (C) and (D), Chain length distribution of the wild-type amylopectin purified from the CL2B gel permeation chromatography from exponentially growing cells (C) and cells grown five days under nitrogen starvation (D). The results are the average CLD of three independent experiments performed on both mutant alleles and are displayed as the molar percentage of each chain. The continuous lines correspond to the difference plot determined for each glucan between the average distribution observed in the wild type and the average distribution obtained from the mutant amylopectins (i.e., mutant profile minus wild-type profile; *be1* mutants in grey, *be2* mutants in red and *be3* mutants in green). Chain lengths (degrees of polymerization) are indicated on the x-axis. The percentage of each class of glucans is presented on the y-axis. The percentages of each chain ± SD for the amylopectin produced by each mutant allele can be found in supplemental figure 5.

We also studied the chain length distribution of amylopectin thanks to the separation of APTS-labeled chains by capillary electrophoresis (Fig. 5). These analyses revealed changes in the chain length composition for the chains composed of 5 to 45 glucose residues. Figures 5c and 5d were obtained by subtracting the average chain length distribution of the amylopectin produced by the different mutant alleles for both transitory (Fig. 5C) and storage starch (Fig. 5D) from the corresponding distribution of the wild-type. The mean values ± SD obtained from three independent chain length distribution experiments of each amylopectin are shown in Fig. S5. The amylopectin molecules produced by *be1* mutants and purified from both transitory and storage starch display a chain length pattern very close to that of the wild type strain grown under the same conditions (Fig. 5C and 5D). For amylopectin purified from transitory starch (Fig. 5c), the lack of BE3 resulted in a significant decrease in chains composed of 6 to 14 glucose residues. The opposite observation was made in *be2* mutants, in which the amylopectin molecule contained an increased amount of short chain (DP 8-13) that was accompanied by a decrease in the amount of chains containing 15 to 55 glucose residues. The same differences were observed between wild-type and *be3* mutant amylopectins for storage starch (Fig. 5D). The amylopectin purified from storage starch produced by *be2* mutants had a chain length distribution resembling that of *be3* mutants, but the decrease in very short chains was restricted to DP 7 to 10 accompanied by an increase of chains with DP 11 to 18 (Fig. 5d). These analyses of chain length distributions allowed us to estimate the branching ratio of each polysaccharide (Szydlowski et al., 2011) (Table 2). These calculated branching levels were all in the range usually described for amylopectin (Table 2). No significant modifications were measured in our mutants with the noticeable exception of the *be3* mutants for which a 20 % decrease in the branching level was observed for both transitory and storage starches.

**Table 2:** Branching levels of wild-type and mutant amylopectins. The branching ratio in amylopectin produced by the wild-type and the mutant strains were determined from the chain length distribution of the corresponding polysaccharides. Branching ratio were calculated with the following equation: 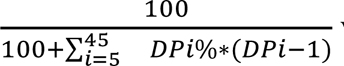 where DPi% represents the percentage of the glucan within the population analyzed and DPi is its length. The results are displayed as means ± SD of calculations from three independent chain length distributions of each amylopectin. Statically significant differences with the wild-type values are indicated by a star (p<0,05).

|  | Transitory<br>starch | Storage<br>starch |
| --- | --- | --- |
| CC4533 | 6.12 ± 0.24 % | 6.66 ± 0.20 % |
| *be1-1* | 6.38 ± 0.12 % | 6.54 ± 0.17 % |
| *be1-2* | 6.29 ± 0.09 % | 6.64 ± 0.07 % |
| *be2-1* | 6.70 ± 0.24 % | 6.58 ± 0.13 % |
| *be2-2* | 6.52 ± 0.17 % | 6.48 ± 0.22 % |
| *be3-1* | 5.74 ± 0.11 %* | 5.78 ± 0.14 %* |
| *be3-3* | 5.80 ± 0.17 %* | 5.36 ± 0.30 %* |

### Determination of the chain length transfer specificities of Chlamydomonas branching enzymes

The differences observed in the Chlamydomonas BE mutants may reflect distinct enzymatic specificities of the corresponding enzymes. To test this hypothesis, the three Chlamydomonas BEs were produced in their recombinant form in *E. coli* (see material and methods; Fig. 6). A high level of purification was obtained for all enzyme preparations as evidenced by SDS-PAGE (Fig. 6A). Zymogram analysis showed no contaminating hydrolytic activities in the enzyme preparations (Fig. 6B). Indeed, a single activity band corresponding to the purified branching enzyme was detected on these activity gels containing starch as a substrate. Different amounts of each purified enzyme were incubated in the presence of amylose in time course experiments in order to define the conditions for which the reaction rates were proportional to the amount of proteins and the duration of the reaction. In these experiments, the activity of the purified enzymes was assessed by the decrease of the absorbance at 660 nm of the amylose/iodine complex (Fig.6C). The reaction rate on amylose was several times higher for BE1 compared to the two other BE isoforms, suggesting that this polysaccharide is a bad substrate for BE2 and BE3 (Fig. 6C).

**Figure 6:**
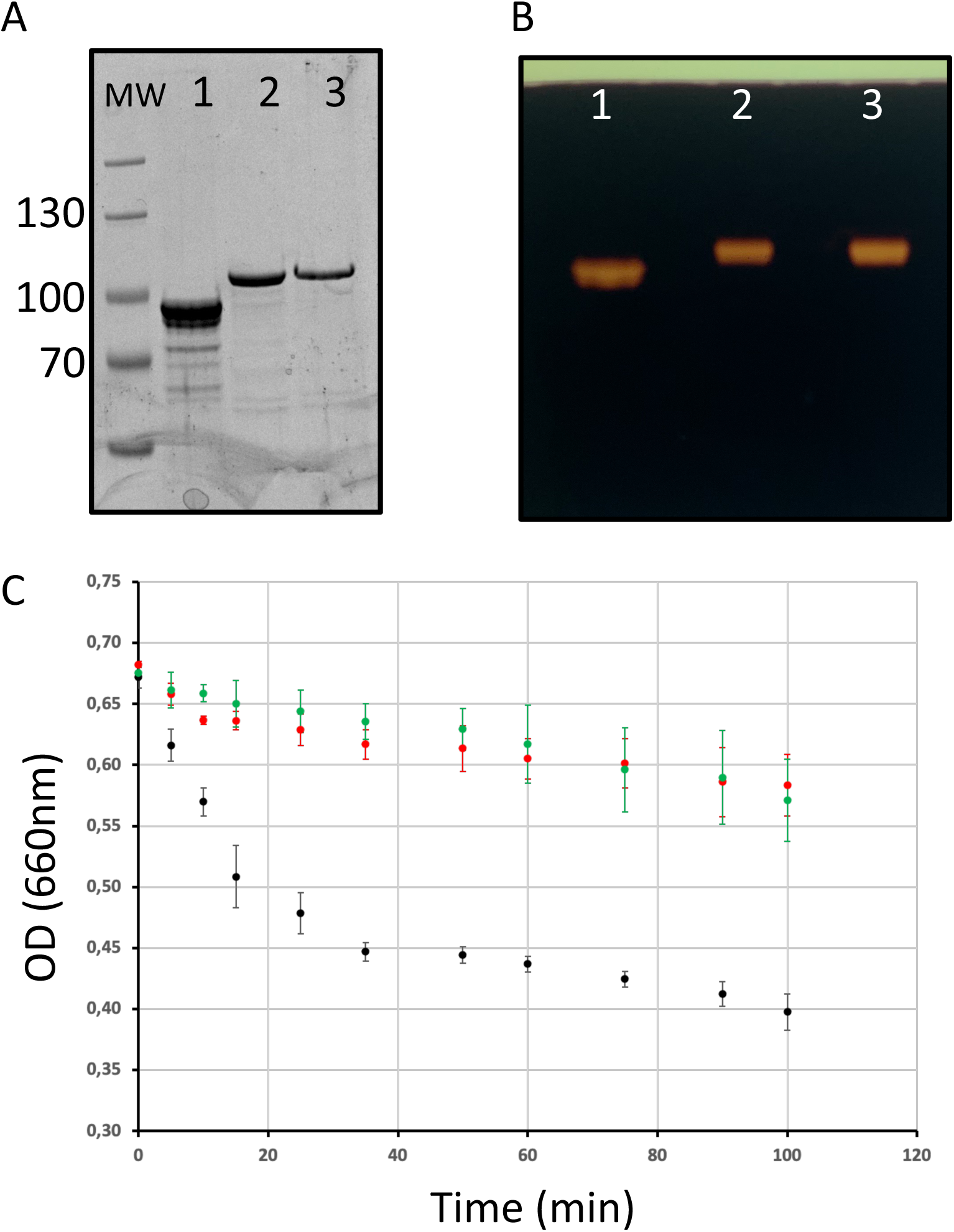
Purification of the recombinant branching enzyme isoforms. (a) SDS-PAGE of the His-tagged recombinant Chlamydomonas starch branching enzyme isoforms. Ten µl of each purified enzymes were analyzed on an 8 % acrylamide SDS-PAGE which was stained with InstantBlue (A) or on a starch-containing zymogram performed in denaturing conditions and stained with iodine after enzyme renaturation (B). Lane 1: 6his-BE1; lane 2: 6-His BE2; lane 3: 6-His BE3; MW: Thermoscientific pre-stained Pageruler. (C) Kinetics of the branching activity of 3 µg of each recombinant BE on potato amylose were evaluated by the decrease in absorbance of the polysaccharide/iodine complex at 660 nm. The results are displayed as the average ± SD of three independent experiments performed with three independent enzyme preparations. The curves obtained with BE1, BE2, and BE3 are displayed in black, red, and green respectively.

To determine the specificities of each BE in terms of length of the transferred chains, each enzyme was incubated with either potato amylopectin or amylose and the newly formed branched chains were characterized after debranching and separation of the APTS-labeled chains by capillary electrophoresis (see material and methods; Fig. 7). For amylose, the results are displayed as the total molar percentages of the chains detected after incubation (Fig. 7A to C). For amylopectin, polysaccharide chain length distributions obtained after incubation were subtracted from that of the original substrate and are presented as a molar percentage since it shows to what extent each chain was increased or decreased upon incubation with the enzyme preparation (Fig. 7D to F). The incubation of amylose with the recombinant BE1 led to the formation of a wide range of newly branched chains with sizes ranging from DP 6 to DP 50. Branched chains with sizes comprised between 6 and 11 glucose residues accounted for more than 50 % of the total chains created (Fig. 7A). Conversely, both BE2 and BE3 created ramified amylose with a narrower size distribution and displayed high specificities for the lengths of the transferred chains (Fig. 7B and 7C). The newly formed chain lengths were ranging from DP 6 to 25 and DP 6 to 16 for BE2 and BE3 respectively (Fig. 7B and 7C). BE3 showed a high level of specificity, with more than 80 % of the total number of newly created branches being exclusively composed of 6 or 7 glucose residues (Fig. 7C). The transfer reaction mediated by BE2 appeared to be less selective with new chains ranging from DP 6 to DP 14. Nevertheless, the chains containing 6 to 9 residues still accounted for more than 70 % of the total newly branched chains (Fig. 7B). These differences were also measured when the purified recombinant enzymes were incubated with potato amylopectin (Fig. 7D to F). Both incubations with BE2 or BE3 led to a marked increase in chains with DP 6 and DP 7 and to a lesser extent in chains with DP 8 to DP 10 for BE2 (Fig. 7E and 7F). Chains with DP 11 to 32 and DP 10 to 30 decreased after the action of BE2 and BE3 respectively. The pattern obtained after the enzymatic reaction of BE1 with amylopectin was clearly different from that of BE2 or BE3 (Fig. 7D) with the observation of two peaks corresponding respectively to short chains (with DP 6 to 12) and to chains sizes ranging from 27 to 36 glucose residues (Fig. 7D). The modified amylopectin molecules display a decrease for chains of DP 13 to 26 and DP > 39. Finally, while a discrete peak of DP 31 to 35 glucans was systematically observed upon incubation of BE2 with amylopectin, we failed to detect any chains with DP over 9 in the reaction products of BE3 with amylopectin.

**Figure 7:**
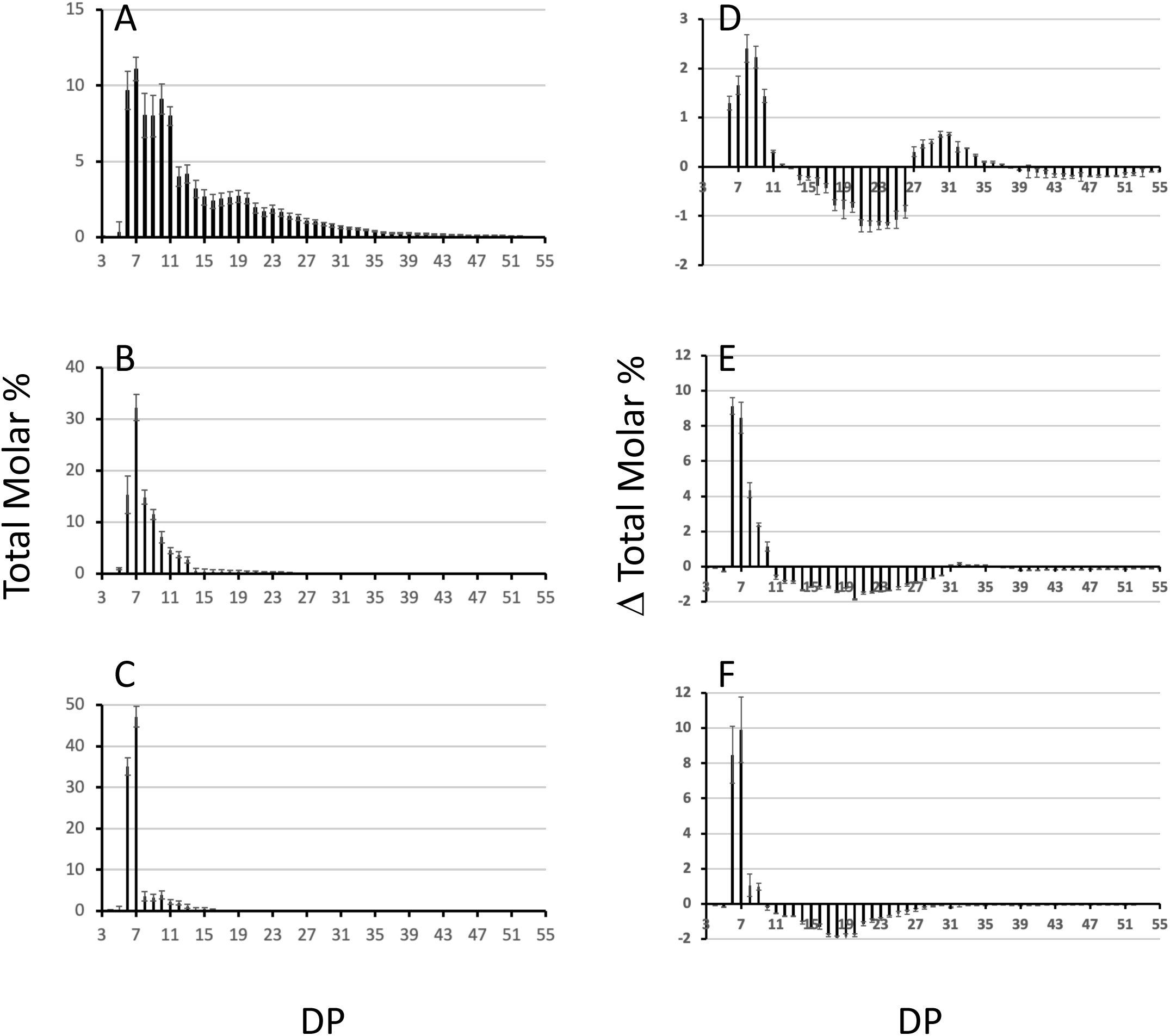
Chain length distribution of the glucans produced by the Chlamydomonas recombinant BEs. One unit of each purified enzyme was incubated in the presence of potato amylose (a to c) or potato amylopectin (d to f) at 30°C for 30 min. The graphs display the average chain length distribution ± SD of three independent experiments of the reaction products obtained upon incubation with BE1 (A and D), BE2 (B and E), and BE3 (C and F). The graphs a to c correspond to the total molar percentages of the chains detected after enzymatic debranching of the reaction products obtained from amylose while graphs d to f display the differential plots obtained by subtracting the CLD of the reaction products to the CLD of unmodified potato amylopectin. Degrees of polymerization (DP) are indicated on the x-axis.

## Discussion

In this study, we sought to determine the impact of the individual loss of each branching enzyme isoform on starch synthesis in Chlamydomonas. The genome of Chlamydomonas contains three isoforms of starch branching enzymes annotated BE1, BE2 and BE3 respectively. The three Chlamydomonas proteins display a typical organization and 3D-structure of starch branching enzyme including the Asp-Glu-Asp catalytic triad whose significance was determined by site-mutagenesis studies with a recombinant maize enzyme (Binderup et al., 2002; Kuriki et al., 1996), and two conserved histidine residues critical for substrate binding (Funane et al., 1998) (Fig. 1B). Sequence alignments, gene phylogeny and protein structure predictions all argue in favor of a proximity between Chlamydomonas BE1 with land plants type 1 BEs. Additionally, BE2 and BE3 are Chlamydomonadales specific paralogs that share ancestry as well as several features with type 2 BEs of Embryophytes. In particular, we could determine a close structural similarity between the Chlamydomonas isoforms and the rice branching enzymes concerning two specific loops which were described as defining the enzyme’s specificity for the length of the transferred chains (Gavgani et al., 2022). All these data point out the existence of a type 1 branching enzyme (BE1) and two type 2 BEs (BE2 and BE3) in Chlamydomonas.

We then analyzed the consequences of the loss of each isoform of BE in Chlamydomonas. The BE1 isoform was detected as a unique activity band on zymograms performed in native conditions while multiple band activities were attributed to both BE2 and BE3 suggesting that these two last isoforms may be part of different protein complexes (Fig. 3). The strains defective at the *BE1* locus displayed no phenotype in the conditions tested in this work. Indeed, these strains accumulate the same starch amounts as the wild-type reference and the polysaccharide produced display the same structural characteristics (Table 1; Fig. 4 and 5). The loss of SBEI in many plant species also failed to reveal any phenotype on starch metabolism, questioning its role in this metabolic pathway. However, subtle modifications in starch structure have been described in a few mutants of land plants. This is the case in maize where the loss of BE1 resulted in an impaired seedling growth (Xia et al., 2011), while the homologous mutant in rice produced a polysaccharide with modified physicochemical properties (Satoh et al., 2003). Interestingly, we isolated a few years ago another *be1* mutant allele which was obtained through the use of a screen aiming at identifying defects in starch degradation (Tunçay et al., 2013) suggesting that the polysaccharide produced by the *be1* mutant is less sensitive to hydrolytic activities. However, we were unable to detect such modifications in the architecture of the polysaccharide produced in our mutants.

Under nitrogen starvation, the Chlamydomonas *be3* strains display a phenotype close to the one reported for land plants *amylose extender* mutants (Banks et al., 1974; Hilbert and MacMasters, 1946; Klucinec and Thompson, 2002; Liu et al., 2011). We, indeed, observed a decrease in the amount of starch produced by these strains (Table 1), as well as an enrichment in amylose, an increase in amylopectin long chains (Table 1 and Figs. 4 and 5), a reduced branch point frequency (Table 2) and a modified chain length distribution of amylopectin with a noticeable reduction for chains with DP comprised between 6 and 15 combined to an increase of chains with DP up to 35 compared with the wild-type (Fig. 5). Transitory starch produced by the *be3* mutants displayed the same differences compared to the wild-type polysaccharide, though, in these growth conditions, those differences were less pronounced. All these observations argue in favor of a major role of the BE3 isoform in both transitory and storage starch syntheses. *be2* mutants display a surprising and opposite situation: storage starch accumulated by these strains contain less amylose and the value of the amylopectin λmax is decreased suggesting a decrease in longer chains ratios of amylopectin, which is indeed confirmed by the analysis of the debranched polysaccharide on TSK-HW50 column (Table 1 and Fig. 4 and 5). Nevertheless, a slight decrease in the short chains of amylopectin (DP < 10) was also observed in this mutant (Fig. 5D), the only phenotype which is expected for a branching enzyme defect. Transitory starch produced by the *be2* mutants displayed an even more unexpected phenotype, with a slight enrichment of amylose and a statistically significant increase in the amount of total starch accumulated (Table 1). Moreover, the amylopectin λmax is really low (555 nm compared to the 580 nm recorded in all the other strains used in this study) which is in the range of the values usually recorded for the wild-type amylopectin found in storage starch (Table 1). Moreover, a decrease in the amylose content and in the long chains of amylopectin was confirmed by the analysis of debranched starch onto gel permeation chromatography (Fig. 5A). Finally, CLD analysis revealed an increase in short chains (DP 8-13) and a slight decrease of longer chains (DP 15-45) in the *be2* mutant amylopectin (Fig. 5c).

The architecture and composition of the polysaccharide produced in our mutants suggest partial redundancy between the activity of BE2 and BE3. BE2 doesn’t seem to be able to compensate for the loss of BE3 since the *be3* mutant strains still display an *amylose extender* phenotype. The situation is reversed in the case of the *be2* mutants in which BE3 not only appears to fully compensate for the loss of BE2 but also leads to an increased production of starch which contains a lower amount of amylose. It is worth noting that similar levels of branching were recorded for both *be2* and wild-type amylopectin. The unexpected phenotypes recorded in the *be2* mutants argue then in favor of an overcompensation phenomenon by the BE3 isoform in Chlamydomonas when BE2 is lacking.

Another explanation for these huge differences in phenotypes between *be2* and *be3* mutants could reside in the respective enzyme substrate affinities and in the length of the chain they transfer. To test this hypothesis, we assayed the three enzymes produced in a recombinant form. First, the BE1 isoform was several times more active on amylose than the two other isoforms, a characteristic of land plants type 1 branching enzymes. The kinetics of appearance of branched chains on amylose were of the same order of magnitude for both recombinant BE2 and BE3 (Fig. 6C). The incubation of BE1 with amylose produced a variety of newly branched glucan lengths ranging from DP 6 to DP 50 with an overrepresentation of short chains (DP 6 to 11). The chains produced by BE2 or BE3 were only composed of short chains with sizes ranging from 6 to 15 glucose residues, even if small differences were detectable between these two isoforms. These small differences were also observed when these enzymes were incubated with amylopectin and confirmed that both BE2 and BE3 isoforms mainly transfer very short chains. In conclusion, both Chlamydomonas BE2 and BE3 display a close proximity with the reaction pattern described for type 2 branching enzymes of land plants (Guan et al., 1997; Nakamura et al., 2010). The small differences assessed between the enzymatic properties of BE2 and BE3 isoforms can hardly explain the totally different phenotypes recorded in the corresponding mutants. Several hypotheses could explain the phenotype differences between our mutants. First, land plant starch BEs are regulated by protein phosphorylation. Experiments performed in wheat and maize revealed that BE were catalytically activated by phosphorylation at one or more serine residues and *in vitro* dephosphorylation led to a reduced activity (Liu et al., 2009; Tetlow et al., 2004, 2008). While the Chlamydomonas BE2 isoform contains all three serine residues involved in the phosphorylation of BEs, only two of them are found in BE3 suggesting that this isoform may be regulated differently. Moreover, phosphorylation of land plant type 2 BEs were shown to be required for the formation of heteromeric protein complexes with other starch metabolism enzymes including starch synthase isoforms, starch phosphorylase or BE1 isoform (Liu et al., 2009, 2012; Tetlow et al., 2004; Subasinghe et al., 2014) . Since we detected both BE2 and BE3 as part of at least two different protein complexes, the absence of one of these isoforms could also affect other enzymes of starch metabolism leading to the production of polysaccharides with modified composition.

In conclusion, our work demonstrates the involvement of two of the three isoforms found in Chlamydomonas in starch synthesis, both displaying characteristics of type 2 branching enzymes. The phenotype of the corresponding mutants suggests a preponderant role of BE3 in the metabolic pathway, this isoform being able to fully compensate for the lack of BE2. In both growth conditions tested, the strains defective for the BE3 isoform displayed the strongest phenotype which is close to the one described for the *amylose extender* mutants of land plants. However, a complete understanding of the precise role of each Chlamydomonas BE isoforms still require an in-depth characterization of the composition of the protein complexes detected in both the wild-type and the different *be* mutants. Moreover, in order to estimate the levels of functional redundancy between the BE isoforms, the phenotype of double mutant strains defective for two isoforms would be required. We attempted several times to produce such double mutant strains by crossing but we were not able to obtain any meiotic progeny with the CLiP mutant strains used in this study. However, these double mutant strains could be obtained thanks to CRISPR gene editing, which is now well-optimized in Chlamydomonas (Dhokane et al., 2020; Ghribi et al., 2020; Kelterborn et al., 2022).

## Methods

### Chlamydomonas strains, growth, and media

The starch branching enzyme mutant strains together with their background strain CC5325 used in this study were obtained from the CLiP collection (Li et al., 2016). For each isoform, two independent insertional mutants were selected: LMJ.RY0402.248984 (*be1-1*), LMJ.RY0402.171898 (*be1-2*), LMJ.SG0182.002820 (*be2-1*), LMJ.RY0402.062087 (*be2-2*), LMJ.RY0402.058754 (*be3-1*), and LMJ.RY0402.131761 (*be3-3*). The initially selected *be3-2* mutant strain (LMJ.RY0402.071404) wasn’t containing any insertion at the *BE3* locus and was then removed from this study. All experiments were carried out under continuous light (40 μmol m^−2^ s^−1^) in standard Tris Acetate Phosphate (TAP) medium (Harris, 1989) at 22°C in liquid cultures under vigorous shaking as described in Delrue et al. (1992). Late-log phase cultures were inoculated at a density of 10^4^ cells/mL and harvested at a final density of 2-3.10^6^ cells/mL. For experiments involving nitrogen starvation, cultures were inoculated at 5.10^5^ cells/mL and harvested after 5 days at a final density of 1-2.10^6^ cells/mL. Cell densities in the cultures were determined using a Multisizer 4 Coulter counter (Beckman, Brea, U.S.A).

### Determination of starch levels, purification, gel permeation chromatography of polysaccharides and chain length distribution (CLD)

Starch purification, assays through amyloglucosidase digestion, and λmax (maximum wavelength of the iodine polysaccharide complex) measurements were carried out as described before (Delrue et al., 1992). Separation of amylose and amylopectin by gel permeation chromatography on a Sepharose CL-2B column and amylose ratio determination was performed with the procedure described in Wattebled et al. (2003). Prior to chain length distribution (CLD) analysis or gel permeation chromatography performed on TSK-HW-50 (Merck, Darmstadt, Germany), the polysaccharides were enzymatically debranched with a cocktail of 1.5 unit of *Pseudomonas sp.* isoamylase (Megazyme, Bray, Ireland) and 1 unit of *Klebsiella planticola* pullulanase (Megazyme, Bray, Ireland) for 16 hours at 42°C in 55 mM Sodium Acetate buffer pH 4.5. Linear glucans resulting from the digestion of 10 mg of starch were separated on a TSK-HW-50 column (95 cm long, 1,6 cm inner diameter) eluted in 10% DMSO at a flow rate of 12 ml/h. Eighty microliters of each 2 mL fraction collected were mixed with 20 µl of Lugol (5% KI, 0,5% I2) to monitor the absorbance of the iodine/polysaccharide complexes and their wavelength at maximal absorbance (λmax). The λmax values allowed us to estimate the average chain-length contained in each fraction (Banks et al., 1971). The chain length distribution of wild-type and mutant amylopectin were obtained through separation of the 8-amino-1,3,6-pyrenetrisulfonic acid (APTS)-labelled chains following the procedure fully described in Fermont et al. (2022).

### Protein sequence comparison, structural model inference

Sequence alignment and homology searches of the Chlamydomonas starch branching enzyme isoforms and their land plants homologues were carried out using the multalin online tool available at https://npsa-prabi.ibcp.fr/cgi-bin/npsa_automat.pl?page=/NPSA/npsa_multalin.html and at the Phytozome website (Goodstein et al., 2012). The putative transit peptide sequences and cleavage sites were obtained with the ChloroP prediction software run locally (Emanuelsson et al., 1999). The molecular protein models were computed using the AlphaFold2 algorithm (Jumper et al., 2021) and were superimposed and visualized using Pymol (DeLano, 2017).

### Phylogenetic tree reconstruction

Protein sequences homologous to Chlamydomonas starch branching enzymes were collected by similarity searches using blastp (Altschul et al., 1990) recursively against RefSeq (O’Leary et al., 2016) and against a custom protein sequences database mainly composed of algal proteomes. All hits were combined, dereplicated, and sequence headers were standardized. A rough phylogenetic tree was reconstructed with the following procedure: (1) sequences were aligned using mafft:FFT-NS-2 v7.503 (Katoh and Standley, 2013). The multiple alignment was trimmed using trimal:gappyout v1.2rev59 and used to infer an approximate Maximum Likelihood phylogenetic tree with fasttree:LG+Ⲅ v2.1.11 (Price et al., 2010). The tree and the alignment were manually inspected to exclude (1) sequences not related to Viridiplantae or Rhodophyceae, (2) sequences composing orthologous groups not related to BE, (3) partial sequences. Additionally, the general species diversity was reduced for the sake of readability. The resulting reduced sequences set was realigned using mafft:FFT-NS-2, trimmed with trimal:gappyout, and the final alignment was used for tree reconstruction with iqtree:LG+C20+Ⲅ v1.6.11 including 1000 ultrafast-bootstrap replicates (Nguyen et al., 2014). The resulting consensus tree was rooted using Rhodophyceae as an outgroup and manually decorated for presentation.

### Molecular techniques, recombinant proteins purification

Oligonucleotides used in this study are described and listed in Supplementary Table S1 and were purchased from Eurofins Genomics (Ebersberg, Germany). Genomic DNA and total RNA were extracted from Chlamydomonas cells following established protocols (Rochaix JD, Mayfield S, Goldschmidt-Clermont M, Erickson J., 1991; Merendino et al., 2003). Amplifications were performed on 1μg of genomic DNA (GoTaq polymerase, Promega) or 500 ng of total RNA for reverse transcription using the One Step RT PCR-Kit (Qiagen) following the manufacturer’s instructions. The extension time was set to 1 min per kb to be amplified. For RT-PCR experiments, the reactional mix contained two specific primers annealing at the 3’ end of the gene of interest and two primers annealing to the *PHOB* sequence (the structural gene encoding one of the starch phosphorylase isoforms in Chlamydomonas, *Cre12.g552200*) which was used as an internal control. The complete cDNA encoding the complete starch branching enzyme mature isoforms (devoid of their putative transit peptides) were amplified from a Chlamydomonas cDNA bank using the proof-reading Kapa Hifi Hot start polymerase (Roche Diagnostics, Austria) and cloned into the pENT-D-Topo plasmid (Thermofisher, Rochester,U.S.A). The entry vectors were fully sequenced to ascertain the integrity of the cloned sequences and were used to transfer the branching enzyme sequences to the Champion^TM^ pET300 plasmid (Thermofisher) generating plasmids in which the mature Chlamydomonas proteins are expressed fused to a N-terminal 6-Histidine tag. The recombinant proteins were expressed from 200 mL of *E. coli* Rosetta^TM^ (Novagen) cultures in Luria Broth medium supplemented with 100 µg/mL ampicillin and 20 µg/mL chloramphenicol then induced with 0,5 mM Isopropyl β-D-1-thiogalactopyranoside (IPTG) at 37°C for 3 hours. Cells were pelleted at 5000 g for 20 min at 4°C and immediately stored at −80°C until use. All the purification steps were carried out at 4°C. The frozen cells pellets were resuspended in 20 mL of 20 mM Tris/HCl, 0.3M NaCl, 50mM Imidazole buffer containing DNAse 1 (1µg/mL), disrupted by sonication and centrifuged at 10,000g for 30 min at 4°C. The supernatant was filtered through a 0.22 µm filter then applied to a 5 mL His-Trap^TM^ HP (Sigma) column pre-equilibrated with the same buffer at a flow rate of 1.5 mL.min^-1^. The recombinant proteins were eluted using 50 mM imidazole gradient steps; an aliquot (10 µL) of each 5-mL fraction was analyzed on SDS-PAGE to assess the level of purification. The recombinant BE1, BE2 and BE3 without any detected protein contamination were obtained at concentrations of 200 mM, 150 mM and 200 mM imidazole respectively. The peak fraction of each BE protein was desalted on PD10 desalting column (Sigma) equilibrated in 20 mM Tris/HCl pH 7, 150 mM NaCl, 10% glycerol. The purification level of each recombinant enzyme was estimated by analyzing an aliquot (20 µl) of each purification fraction on SDS-PAGE (8% acrylamide) stained with Instantblue^TM^ (Abcam).

### Zymograms and anion exchange chromatography

Zymograms in starch containing gels allowing the detection of most starch hydrolases have been performed as previously described (Mouille et al., 1996). The semi-purification of the branching enzyme activities was performed from 1 liter of exponential phase TAP cultures of the wild-type strain CC5325 and the characterized BE mutant strains. Algae were spun down at 3000 g for 10 min at 4°C and the cell pellet was disrupted by sonication in 20 mL of prechilled purification buffer (50 mM 1,3-Bis[tris(hydroxymethyl)methylamino]propane/HCl pH 7). The lysate was centrifuged at 10 000 g for 20 min at 4°C and the supernatant was filtered through a 0.22 µm filter then applied to an anion exchange chromatography column (HiTrap^TM^ Q FF, Cytiva) pre-equilibrated in the same buffer at a flow rate of 2mL/min until UV absorption monitored at 280 nm was back to the baseline. Elution of branching enzyme activities was performed in 1 mL fractions by applying a 90 min linear gradient (0-300 mM NaCl in purification buffer) at a flow rate of 1 mL/min. Thirty microliters of each even fraction from fraction 10 to 70 was analyzed on native zymogram based on phosphorylase A stimulation. Proteins were separated at 4°C and 15 V/cm for 3 h onto a native PAGE (6 % acrylamide). After electrophoresis, the gels were washed 20 min in 20 mL of 50 mM Hepes/NaOH pH 7, 10 % glycerol buffer then incubated overnight in the same buffer containing 50 mM glucose-1-phosphate, 2.5 mM AMP, and 28 units of phosphorylase a from rabbit muscle. Branching activities were revealed by iodine staining.

### Branching enzyme assay and analysis of their reaction products

The enzyme activity was measured at 30 °C and pH 7 by monitoring the decrease of the absorbance of the glucan-iodine complex resulting from the branching of potato amylose (Boyer and Preiss, 1978). The purified Chlamydomonas recombinant BE isoforms were incubated at 30°C in the presence of potato amylose (2mg/mL) in 560 µl of 50 mM Tris/HCl pH 7. Changes in the absorbance at 660 nm of the glucan/iodine complex were regularly monitored by mixing a diluted aliquot (40 µl of the reaction mixture in 600 µl of Tris buffer) of the reaction mixture with 160 µl of Lugol reagent (1% KI and 0.1% I2). One enzyme unit was defined as the decrease of 1 unit of absorbance at 660 nm per min at 30°C.

To determine the specificities of each Chlamydomonas BE isoforms, one unit of the corresponding purified recombinant protein was incubated in 500 µl of 50 mM Tris/HCl pH7 buffer containing either potato amylose or potato amylopectin at a final concentration of 2mg/mL for 30 min. The reaction was stopped by boiling the samples for 5 min and the chain length distribution of the modified polysaccharides were analyzed after enzymatic debranching and analysis on capillary electrophoresis following the protocol detailed previously for amylopectin CLD analysis.

## Authors Contribution

AC, OG and TD performed experiments. PD performed the phylogenetic analyses and CB the protein structure prediction with Alphafold 2. PD and CB edited the manuscript. DD designed and performed experiments, and wrote the manuscript. All authors gave final approval for submission.

## Supplemental materials

**Fig. S1**. Structural predictions of *Chlamydomonas reinhardtii* BE isoforms.

**Fig. S2.** Maximum likelihood consensus phylogenetic tree of starch branching enzymes in algae and plants.

**Fig. S3**. Molecular characterization of the *be* mutant strains.

**Fig. S4** Growth curves of Chlamydomonas wild type and *be* mutant strains.

**Fig. S5** Chain length distribution of wild-type and mutant amylopectins..

**Table S1:** List of the primers used in this study.

## References

Altschul, S.F., Gish, W., Miller, W., Myers, E.W., and Lipman, D.J. (1990). Basic local alignment search tool. J. Mol. Biol. 215: 403–410.

Ball, S., Guan, H.P., James, M., Myers, A., Keeling, P., Mouille, G., Buléon, A., Colonna, P., and Preiss, J. (1996). From glycogen to amylopectin: a model for the biogenesis of the plant starch granule. Cell 86: 349–352.

Banks, W., Greenwood, C.T., and Khan, K.M. (1971). The interaction of linear amylose oligomers with iodine. Carbohydr. Res. 17(1): 25–33.

Banks, W., Greenwood, C.T., and Muir, D.D. (1974). Studies on Starches of High Amylose Content. Part 17. A Review of Current Concepts. Starch - Stärke 26: 289–300.

Bhattacharyya, M.K., Smith, A.M., Ellis, T.H., Hedley, C., and Martin, C. (1990). The wrinkled-seed character of pea described by Mendel is caused by a transposon-like insertion in a gene encoding starch-branching enzyme. Cell 60: 115–122.

Binderup, K., Mikkelsen, R., and Preiss, J. (2002). Truncation of the amino terminus of branching enzyme changes its chain transfer pattern. Arch. Biochem. Biophys. 397: 279–285.

Boyer, C.D. and Preiss, J. (1978). Multiple forms of (1 → 4)-α-d-glucan, (1 → 4)-α-d-glucan-6-glycosyl transferase from developing zea mays L. Kernels. Carbohydr. Res. 61: 321–334.

Buléon, A., Gallant, D.J., Bouchet, B., Mouille, G., D’Hulst, C., Kossmann, J., and Ball, S. (1997). Starches from A to C. Chlamydomonas reinhardtii as a model microbial system to investigate the biosynthesis of the plant amylopectin crystal. Plant Physiol. 115: 949–957.

DeLano, W.L. (2017). The PyMOL molecular graphics system (Schrödinger, LLC).

Delrue, B. et al. (1992). Waxy Chlamydomonas reinhardtii: monocellular algal mutants defective in amylose biosynthesis and granule-bound starch synthase activity accumulate a structurally modified amylopectin. J. Bacteriol. 174: 3612–3620.

Deschamps, P., Colleoni, C., Nakamura, Y., Suzuki, E., Putaux, J.-L., Buléon, A., Haebel, S., Ritte, G., Steup, M., Falcón, L.I., and Others (2008). Metabolic symbiosis and the birth of the plant kingdom. Mol. Biol. Evol. 25: 536–548.

Dhokane, D., Bhadra, B., and Dasgupta, S. (2020). CRISPR based targeted genome editing of Chlamydomonas reinhardtii using programmed Cas9-gRNA ribonucleoprotein. Mol. Biol. Rep. 47: 8747–8755.

Emanuelsson, O., Nielsen, H., and von Heijne, G. (1999). ChloroP, a neural network-based method for predicting chloroplast transit peptides and their cleavage sites. Protein Sci. 8: 978–984.

Fermont, L., Szydlowski, N., and Colleoni, C. (2022). Determination of Glucan Chain Length Distribution of Glycogen Using the Fluorophore-Assisted Carbohydrate Electrophoresis (FACE) Method. J. Vis. Exp.

Findinier, J., Laurent, S., Duchêne, T., Roussel, X., Lancelon-Pin, C., Cuiné, S., Putaux, J.-L., Li-Beisson, Y., D’Hulst, C., Wattebled, F., and Dauvillée, D. (2019). Deletion of BSG1 in Chlamydomonas reinhardtii leads to abnormal starch granule size and morphology. Sci. Rep. 9: 1990.

Fontaine, T., D’Hulst, C., Maddelein, M.L., Routier, F., Pépin, T.M., Decq, A., Wieruszeski, J.M., Delrue, B., Van den Koornhuyse, N., and Bossu, J.P. (1993). Toward an understanding of the biogenesis of the starch granule. Evidence that Chlamydomonas soluble starch synthase II controls the synthesis of intermediate size glucans of amylopectin. J. Biol. Chem. 268: 16223– 16230.

Funane, K., Libessart, N., Stewart, D., Michishita, T., and Preiss, J. (1998). Analysis of essential histidine residues of maize branching enzymes by chemical modification and site-directed mutagenesis. J. Protein Chem. 17: 579–590.

Gao, M., Fisher, D.K., Kim, K.N., Shannon, J.C., and Guiltinan, M.J. (1997). Independent genetic control of maize starch-branching enzymes IIa and IIb. Isolation and characterization of a Sbe2a cDNA. Plant Physiol. 114: 69–78.

Gavgani, H.N., Fawaz, R., Ehyaei, N., Walls, D., Pawlowski, K., Fulgos, R., Park, S., Assar, Z., Ghanbarpour, A., and Geiger, J.H. (2022). A structural explanation for the mechanism and specificity of plant branching enzymes I and IIb. J. Biol. Chem. 298: 101395.

Ghribi, M., Nouemssi, S.B., Meddeb-Mouelhi, F., and Desgagné-Penix, I. (2020). Genome Editing by CRISPR-Cas: A Game Change in the Genetic Manipulation of Chlamydomonas. Life 10.

Goodstein, D.M., Shu, S., Howson, R., Neupane, R., Hayes, R.D., Fazo, J., Mitros, T., Dirks, W., Hellsten, U., Putnam, N., and Rokhsar, D.S. (2012). Phytozome: a comparative platform for green plant genomics. Nucleic Acids Res. 40: D1178–86.

Guan, H., Li, P., Imparl-Radosevich, J., Preiss, J., and Keeling, P. (1997). Comparing the Properties ofEscherichia coliBranching Enzyme and Maize Branching Enzyme. Arch. Biochem. Biophys. 342: 92–98.

Harris, E.H. (2001). CHLAMYDOMONAS AS A MODEL ORGANISM. Annu. Rev. Plant Physiol. Plant Mol. Biol. 52: 363–406.

Harris, E.H. (1989). Culture and Storage Methods. In The Chlamydomonas Sourcebook (Elsevier), pp. 25–63.

Hedman, K.D. and Boyer, C.D. (1982). Gene dosage at the amylose-extender locus of maize: effects on the levels of starch branching enzymes. Biochem. Genet. 20: 483–492.

Hennen-Bierwagen, T.A., Lin, Q., Grimaud, F., Planchot, V., Keeling, P.L., James, M.G., and Myers, A.M. (2009). Proteins from multiple metabolic pathways associate with starch biosynthetic enzymes in high molecular weight complexes: a model for regulation of carbon allocation in maize amyloplasts. Plant Physiol. 149: 1541–1559.

Hennen-Bierwagen, T.A., Liu, F., Marsh, R.S., Kim, S., Gan, Q., Tetlow, I.J., Emes, M.J., James, M.G., and Myers, A.M. (2008). Starch biosynthetic enzymes from developing maize endosperm associate in multisubunit complexes. Plant Physiol. 146: 1892–1908.

Hicks, G.R., Hironaka, C.M., Dauvillee, D., Funke, R.P., D’Hulst, C., Waffenschmidt, S., and Ball, S.G. (2001). When simpler is better. Unicellular green algae for discovering new genes and functions in carbohydrate metabolism. Plant Physiol. 127: 1334–1338.

Hilbert, G.E. and MacMasters, M.M. (1946). Pea starch, a starch of high amylose content. J. Biol. Chem. 162: 229–238.

Hong, S. and Preiss, J. (2000). Localization of C-terminal domains required for the maximal activity or for determination of substrate preference of maize branching enzymes. Arch. Biochem. Biophys. 378: 349–355.

Jumper, J. et al. (2021). Highly accurate protein structure prediction with AlphaFold. Nature 596: 583–589.

Katoh, K. and Standley, D.M. (2013). MAFFT multiple sequence alignment software version 7: improvements in performance and usability. Mol. Biol. Evol. 30: 772–780.

Kelterborn, S., Boehning, F., Sizova, I., Baidukova, O., Evers, H., and Hegemann, P. (2022). Gene Editing in Green Alga Chlamydomonas reinhardtii via CRISPR-Cas9 Ribonucleoproteins. Methods Mol. Biol. 2379: 45–65.

Klucinec, J.D. and Thompson, D.B. (2002). Structure of amylopectins from ae-containing maize starches. Cereal Chem. 79: 19–23.

Kuriki, T., Guan, H., Sivak, M., and Preiss, J. (1996). Analysis of the active center of branching enzyme II from maize endosperm. J. Protein Chem. 15: 305–313.

Kuriki, T., Stewart, D.C., and Preiss, J. (1997). Construction of chimeric enzymes out of maize endosperm branching enzymes I and II: activity and properties. J. Biol. Chem. 272: 28999– 29004.

Libessart, N., Maddelein, M.L., Koornhuyse, N., Decq, A., Delrue, B., Mouille, G., D’Hulst, C., and Ball, S. (1995). Storage, Photosynthesis, and Growth: The Conditional Nature of Mutations Affecting Starch Synthesis and Structure in Chlamydomonas. Plant Cell 7: 1117–1127.

Liu, F., Ahmed, Z., Lee, E.A., Donner, E., Liu, Q., Ahmed, R., Morell, M.K., Emes, M.J., and Tetlow, I.J. (2011). Allelic variants of the amylose extender mutation of maize demonstrate phenotypic variation in starch structure resulting from modified protein–protein interactions. J. Exp. Bot. 63: 1167–1183.

Liu, F., Makhmoudova, A., Lee, E.A., Wait, R., Emes, M.J., and Tetlow, I.J. (2009). The amylose extender mutant of maize conditions novel protein–protein interactions between starch biosynthetic enzymes in amyloplasts. J. Exp. Bot. 60: 4423–4440.

Liu, F., Romanova, N., Lee, E.A., Ahmed, R., Evans, M., Gilbert, E.P., Morell, M.K., Emes, M.J., and Tetlow, I.J. (2012). Glucan affinity of starch synthase IIa determines binding of starch synthase I and starch-branching enzyme IIb to starch granules. Biochem. J 448: 373–387.

Li, X. et al. (2019). A genome-wide algal mutant library and functional screen identifies genes required for eukaryotic photosynthesis. Nat. Genet. 51: 627–635.

Li, X., Zhang, R., Patena, W., Gang, S.S., Blum, S.R., Ivanova, N., Yue, R., Robertson, J.M., Lefebvre, P.A., Fitz-Gibbon, S.T., Grossman, A.R., and Jonikas, M.C. (2016). An Indexed, Mapped Mutant Library Enables Reverse Genetics Studies of Biological Processes in Chlamydomonas reinhardtii. Plant Cell 28: 367–387.

Lombard, V., Golaconda Ramulu, H., Drula, E., Coutinho, P.M., and Henrissat, B. (2014). The carbohydrate-active enzymes database (CAZy) in 2013. Nucleic Acids Res. 42: D490–5.

Maddelein, M.L., Libessart, N., Bellanger, F., Delrue, B., D’Hulst, C., Van den Koornhuyse, N., Fontaine, T., Wieruszeski, J.M., Decq, A., and Ball, S. (1994). Toward an understanding of the biogenesis of the starch granule. Determination of granule-bound and soluble starch synthase functions in amylopectin synthesis. J. Biol. Chem. 269: 25150–25157.

Manners, D.J. (1989). Recent developments in our understanding of amylopectin structure. Carbohydr. Polym. 11: 87–112.

Merchant, S.S. et al. (2007). The Chlamydomonas genome reveals the evolution of key animal and plant functions. Science 318: 245–250.

Merendino, L., Falciatore, A., and Rochaix, J.-D. (2003). Expression and RNA binding properties of the chloroplast ribosomal protein S1 from Chlamydomonas reinhardtii. Plant Mol. Biol. 53: 371–382.

Mizuno, K., Kawasaki, T., Shimada, H., Satoh, H., Kobayashi, E., Okumura, S., Arai, Y., and Baba, T. (1993). Alteration of the structural properties of starch components by the lack of an isoform of starch branching enzyme in rice seeds. J. Biol. Chem. 268: 19084–19091.

Mouille, G., Maddelein, M.L., Libessart, N., Talaga, P., Decq, A., Delrue, B., and Ball, S. (1996). Preamylopectin Processing: A Mandatory Step for Starch Biosynthesis in Plants. Plant Cell 8: 1353–1366.

Nakamura, Y., Utsumi, Y., Sawada, T., Aihara, S., Utsumi, C., Yoshida, M., and Kitamura, S. (2010). Characterization of the reactions of starch branching enzymes from rice endosperm. Plant Cell Physiol. 51: 776–794.

Nguyen, L.T.L.-T., Schmidt, H. a., von Haeseler, A., and Minh, B.Q. (2014). IQ-TREE: A fast and effective stochastic algorithm for estimating maximum-likelihood phylogenies. Mol. Biol. Evol. 32: 268–274.

O’Leary, N.A. et al. (2016). Reference sequence (RefSeq) database at NCBI: current status, taxonomic expansion, and functional annotation. Nucleic Acids Res. 44: D733–45.

Pérez, S. and Bertoft, E. (2010). The molecular structures of starch components and their contribution to the architecture of starch granules: A comprehensive review. Starke 62: 389–420.

Price, M.N., Dehal, P.S., and Arkin, A.P. (2010). FastTree 2--approximately maximum-likelihood trees for large alignments. PLoS One 5: e9490.

Regina, A., Kosar-Hashemi, B., Li, Z., Rampling, L., Cmiel, M., Gianibelli, M.C., Konik-Rose, C., Larroque, O., Rahman, S., and Morell, M.K. (2004). Multiple isoforms of starch branching enzyme-I in wheat: lack of the major SBE-I isoform does not alter starch phenotype. Funct. Plant Biol. 31: 591–601.

Rochaix JD, Mayfield S, Goldschmidt-Clermont M, Erickson J. (1991). Molecular biology of Chlamydomonas. In Plant molecular biology: a practical approach, C.H. Shaw, ed (Oxford: IRL Press), pp. 253–275.

Satoh, H., Nishi, A., Yamashita, K., Takemoto, Y., Tanaka, Y., Hosaka, Y., Sakurai, A., Fujita, N., and Nakamura, Y. (2003). Starch-branching enzyme I-deficient mutation specifically affects the structure and properties of starch in rice endosperm. Plant Physiol. 133: 1111–1121.

Sestili, F., Janni, M., Doherty, A., Botticella, E., D’Ovidio, R., Masci, S., Jones, H.D., and Lafiandra, D. (2010). Increasing the amylose content of durum wheat through silencing of the SBEIIagenes. BMC Plant Biol. 10: 144.

Stam, M.R., Danchin, E.G.J., Rancurel, C., Coutinho, P.M., and Henrissat, B. (2006). Dividing the large glycoside hydrolase family 13 into subfamilies: towards improved functional annotations of α-amylase-related proteins. Protein Eng. Des. Sel. 19: 555–562.

Subasinghe, R.M., Liu, F., Polack, U.C., Lee, E.A., Emes, M.J., and Tetlow, I.J. (2014). Multimeric states of starch phosphorylase determine protein–protein interactions with starch biosynthetic enzymes in amyloplasts. Plant Physiol. Biochem. 83: 168–179.

Sun, C., Sathish, P., Ahlandsberg, S., and Jansson, C. (1998). The two genes encoding starch-branching enzymes IIa and IIb are differentially expressed in barley. Plant Physiol. 118: 37–49.

Szydlowski, N., Ragel, P., Hennen-Bierwagen, T.A., Planchot, V., Myers, A.M., Mérida, A., d’Hulst, C., and Wattebled, F. (2011). Integrated functions among multiple starch synthases determine both amylopectin chain length and branch linkage location in Arabidopsis leaf starch. J. Exp. Bot. 62: 4547–4559.

Tetlow, I.J., Beisel, K.G., Cameron, S., Makhmoudova, A., Liu, F., Bresolin, N.S., Wait, R., Morell, M.K., and Emes, M.J. (2008). Analysis of protein complexes in wheat amyloplasts reveals functional interactions among starch biosynthetic enzymes. Plant Physiol. 146: 1878– 1891.

Tetlow, I.J., Wait, R., Lu, Z., Akkasaeng, R., Bowsher, C.G., Esposito, S., Kosar-Hashemi, B., Morell, M.K., and Emes, M.J. (2004). Protein phosphorylation in amyloplasts regulates starch branching enzyme activity and protein-protein interactions. Plant Cell 16: 694–708.

Tunçay, H., Findinier, J., Duchêne, T., Cogez, V., Cousin, C., Peltier, G., Ball, S.G., and Dauvillée, D. (2013). A forward genetic approach in Chlamydomonas reinhardtii as a strategy for exploring starch catabolism. PLoS One 8: e74763.

Wattebled, F., Ral, J.-P., Dauvillée, D., Myers, A.M., James, M.G., Schlichting, R., Giersch, C., Ball, S.G., and D’Hulst, C. (2003). STA11, a Chlamydomonas reinhardtii locus required for normal starch granule biogenesis, encodes disproportionating enzyme. Further evidence for a function of alpha-1,4 glucanotransferases during starch granule biosynthesis in green algae. Plant Physiol. 132: 137–145.

Xia, H., Yandeau-Nelson, M., Thompson, D.B., and Guiltinan, M.J. (2011). Deficiency of maize starch-branching enzyme I results in altered starch fine structure, decreased digestibility and reduced coleoptile growth during germination. BMC Plant Biol. 11: 95.

Yamanouchi, H. and Nakamura, Y. (1992). Organ Specificity of Isoforms of Starch Branching Enzyme (Q-Enzyme) in Rice. Plant Cell Physiol. 33: 985–991.

Yandeau-Nelson, M.D., Laurens, L., Shi, Z., Xia, H., Smith, A.M., and Guiltinan, M.J. (2011). Starch-branching enzyme IIa is required for proper diurnal cycling of starch in leaves of maize. Plant Physiol. 156: 479–490.

